# Bcl6 promotes neurogenic conversion through transcriptional repression of multiple self-renewal-promoting extrinsic pathways

**DOI:** 10.1101/370999

**Authors:** Jerome Bonnefont, Luca Tiberi, Jelle van den Ameele, Delphine Potier, Zachary B Gaber, Xionghui Lin, Angéline Bilheu, Adèle Herpoel, Fausto D. Velez Bravo, François Guillemot, Stein Aerts, Pierre Vanderhaeghen

**Affiliations:** Université Libre de Bruxelles (U.L.B.), Institut de Recherches en Biologie Humaine et Moléculaire (IRIBHM), and ULB Neuroscience Institute (UNI), 1070 Brussels, Belgium; VIB-KU Leuven Center for Brain & Disease Research, 3000 Leuven, Belgium; The Francis Crick Institute, London NW1 1AT, United Kingdom; Department of Neurosciences, Leuven Brain Institute, KULeuven, 3000 Leuven, Belgium; Welbio, Université Libre de Bruxelles (U.L.B.), B-1070 Brussels, Belgium

**Keywords:** Bcl6, brain development, neurogenesis, stemness, transcription, Notch signaling, Wnt signaling, SHH signaling, FGF signaling, cyclins

## Abstract

During neurogenesis, progenitors switch from self-renewal to differentiation through the interplay of intrinsic and extrinsic cues, but how these are integrated remains poorly understood. Here we combine whole genome transcriptional and epigenetic analyses with in vivo functional studies and show that Bcl6, a transcriptional repressor known to promote neurogenesis, acts as a key driver of the neurogenic transition through direct silencing of a selective repertoire of genes belonging to multiple extrinsic pathways promoting self-renewal, most strikingly the Wnt pathway. At the molecular level, Bcl6 acts through both generic and pathway-specific mechanisms. Our data identify a molecular logic by which a single cell-intrinsic factor ensures robustness of neural cell fate transition by decreasing responsiveness to the extrinsic pathways that favor self-renewal.

## Introduction

During neural development, the generation of the appropriate type and number of differentiated neurons and glial cells is controlled by a complex interplay between extrinsic and intrinsic cues acting on neural progenitors, which control the balance between differentiation and self-renewal (Martynoga et al., 2012; Rossi et al., 2017; Tiberi et al., 2012b).

In the developing cortex, radial glial cells are the main progenitors that will differentiate into specific post-mitotic neuron populations directly or through various classes of intermediate progenitors (Gotz and Huttner, 2005; Kriegstein and Alvarez-Buylla, 2009). Proneural transcription factors act on these progenitors as the main intrinsic drivers of neurogenesis (Guillemot and Hassan, 2017; Guillemot et al., 2006), by directly inducing various classes of genes involved in neuronal differentiation and through cross-repression with the Notch pathway, which promotes self-renewal. Key features of the Notch signaling pathway including lateral inhibition and oscillatory behaviour, contribute to the irreversible commitment of differentiating cells towards neuronal fate (Kageyama et al., 2008). In addition however, many classes of extrinsic morphogen cues, including Wnts, Sonic Hedgehog (SHH), and Fibroblast Growth Factors (FGF) can act on cortical progenitors to promote expansion and self-renewal, and thereby inhibit neurogenesis (Chenn and Walsh, 2002; Kang et al., 2009; Lien et al., 2006; Rash et al., 2011; Wang et al., 2016).

Intriguingly, it has long been proposed that postmitotic cells ongoing neuronal differentiation become insulated from extrinsic signaling (Edlund and Jessell, 1999), but the molecular mechanisms remain essentially unknown. Whether and how the negative modulation of responsiveness to extrinsic cues is required for neuronal commitment remains currently unclear, as are the underlying mechanisms. Delamination of the progenitors away from the ventricular zone could contribute to this process, as some of the morphogen cues are secreted in the embryonic cerebrospinal fluid and are thought to act through the apical processes or cilia of the radial glial cells (Lehtinen et al., 2011). However several cues, most strikingly Wnts, are also present in the cortical tissue (Harrison-Uy and Pleasure, 2012), where they can act on progenitors to block differentiation.

Moreover, the signaling components of these various pathways, as well as some key downstream targets, are often partially overlapping. For instance, cross-talk of the Notch and Wnt pathways (Hayward et al., 2008), or Notch and FGFs (Rash et al., 2011), has been documented during embryonic development and despite the relatively simple intracellular regulation of the Wnt/β-catenin pathway, many of its components are used by other pathways or participate in distinct cellular activities. For instance, the deletion of *Gsk3a/b*, a major intracellular component of the β-catenin destruction complex, increases the proliferation of radial glial cells at the expense of their differentiation by altering not only Wnt but also Notch and FGF signaling activity (Kim et al., 2009). Also, some key effectors genes such as *Cyclin d1/d2* are found as common targets of most morphogen pathways depending on cellular context (Cohen et al., 2010; Kalita et al., 2013; Katoh and Katoh, 2009; Nilsson et al., 2012; Shtutman et al., 1999). How these intermingled pathways are effectively shut down during neurogenesis is therefore an important and complex issue that remains largely unresolved.

We previously reported that the transcriptional repressor Bcl6 (Baron et al., 1993; Chang et al., 1996) is required for neuronal differentiation in the cerebral cortex at least in part through repression of the Notch-dependent *Hes5* target (Tiberi et al., 2012a), while in the cerebellum, Bcl6 promotes neurogenesis through repression of SHH pathway effectors *Gli1/2* (Tiberi et al., 2014). This raises the question whether Bcl6 promotes neurogenic conversion through the repression of distinct targets depending on the cellular context, or through a more generic transcriptional repression programme. Here we combine transcriptome, epigenome and *in vivo* functional analyses to determine the molecular logic of action of Bcl6 during neurogenesis, focusing on the cerebral cortex. We find that Bcl6 acts a global repressor of a repertoire of signaling components of most signaling pathways known to promote self-renewal, including Notch, SHH, FGFs, and most strikingly the Wnt pathway. These data define a molecular logic of neurogenesis whereby a single intrinsic factor can modulate negatively the responsiveness to several extrinsic cues, through transcriptional repression at multiple parallel and serial levels along their downstream pathways, to ensure irreversible neurogenic fate transition.

## Results

### Bcl6 upregulates an intrinsic neurogenic program and downregulates extrinsic proliferative pathways

To determine the primary molecular mechanisms of Bcl6 action in cortical neurogenesis, we performed RNA-seq transcriptome analysis on *in vitro* embryonic stem cell-derived cortical progenitors driving inducible Bcl6 expression (Figure 1A; (Gaspard et al., 2008; Tiberi et al., 2012a). We found that 24 hours following Bcl6 induction 764 genes were significantly up-regulated, with *Bcl6* being the most increased, and 610 genes were significantly down-regulated, with *Hes5* being the most repressed (Supplementary Table S1).

**Figure 1.**
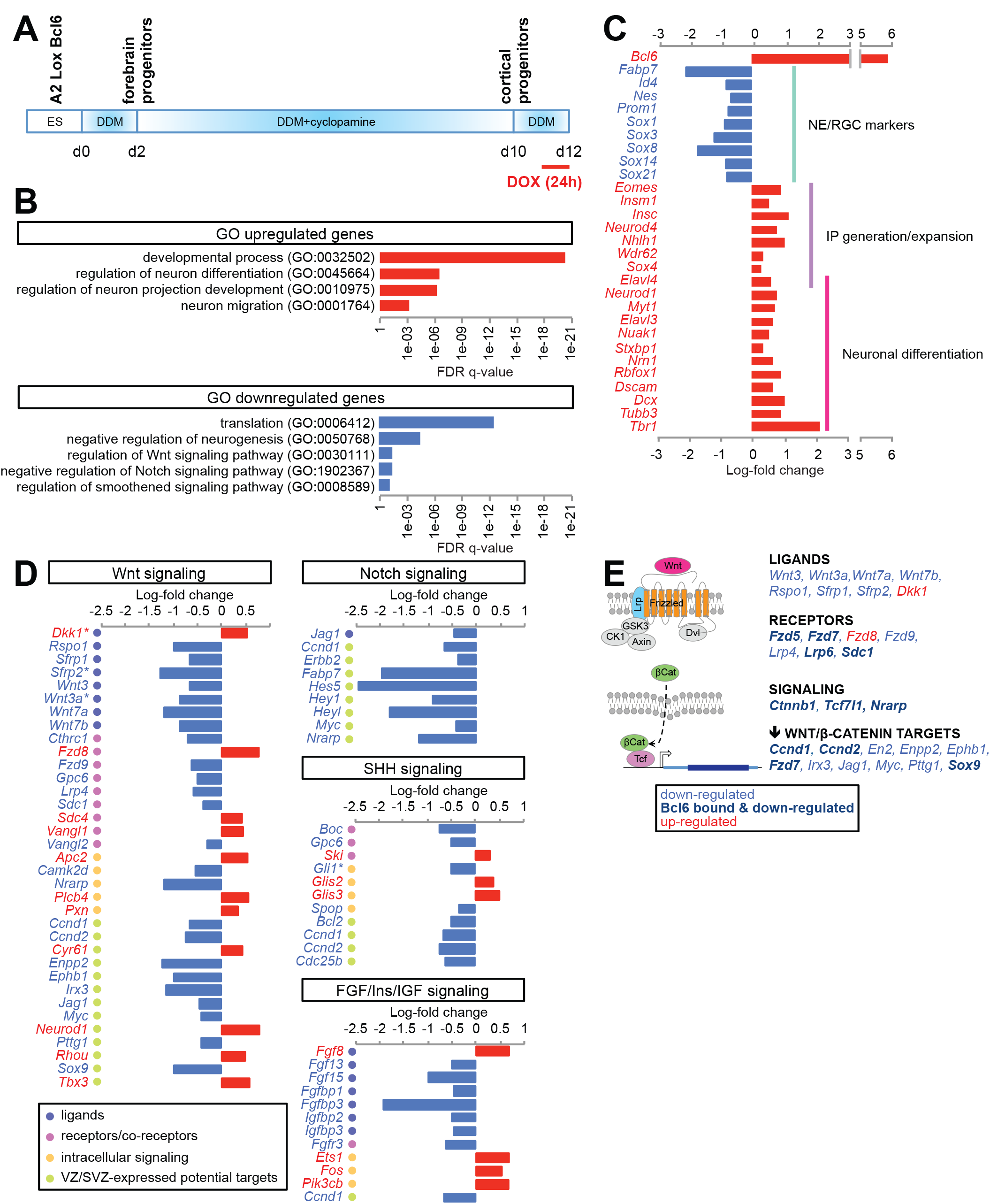
Bcl6 promotes a neurogenic transcription program and represses selective genes of the main proliferative pathways. (A) Scheme representing the differentiation protocol of Bcl6-inducible A2 lox.Cre mouse embryonic stem cells into cortical progenitors. Bcl6 expression was induced at day 12 using a single doxycycline pulse for 24 h. (B) Gene Ontology analysis showing statistically significant enrichment for some categories of up- and down-regulated genes following Bcl6 induction (see also Supplementary Table S2 for complete lists). (C) Histograms representing the log-fold change of a series of significantly up- or down-regulated genes selected upon their expression and/or function during cortical differentiation. NE: neuroepithelial cells, RGC: radial glial cells, IP: intermediate progenitors. (see also Supplementary Table S1 for complete lists). (D) Histograms representing the log-fold change of significantly up- or down-regulated genes belonging to the main proliferative pathways in cortical progenitors. Only the potential target genes with known expression in embryonic cortical progenitors are indicated. For the complete list of genes taken into consideration for the analysis, see also Supplementary Table S3. Genes marked with an asterisk also are target genes of the pathway itself. VZ/SVZ: ventricular and subventricular zones. (E) Scheme of the canonical Wnt pathway depicting the role in the cascade of the ensemble of Wnt/β-catenin-related genes bound and/or altered by Bcl6 investigated in this study.

Gene Ontology analysis of the up-regulated genes revealed a significant enrichment in categories linked to development and cell/neuron differentiation (Figure 1B, Supplementary Table S2). More specifically, most of the canonical markers of differentiation into intermediate progenitors and neurons were significantly up-regulated (Figure 1C;; Supplementary Table S1). On the other hand, down-regulated genes showed an enrichment in gene categories linked to regulation of translation, negative regulation of neurogenesis, and most strikingly to signaling pathways promoting the expansion and self-renewal of cortical progenitors (Figure 1B, Supplementary Table S2). Among the significantly down-regulated genes, we found as expected markers of radial glial cells and transcriptional targets of Notch, but also many signaling components of FGF and SHH-dependent pathways, and most strikingly a high number of genes belonging to the Wnt signaling cascade, from ligands to receptors to target genes (Figures 1C-E; Supplementary Tables S1 and S3). This suggests that Bcl6 could act by repressing multiple proliferative pathways to promote neurogenesis.

### *Bcl6* functionally alters β-catenin signaling to promote neurogenesis

Given the importance of the Wnt pathway in the regulation of self-renewal/differentiation balance in the cortex (Chenn and Walsh, 2002; Fang et al., 2013; Hirabayashi and Gotoh, 2005; Hirabayashi et al., 2004; Kuwahara et al., 2010; Munji et al., 2011; Mutch et al., 2010; Wrobel et al., 2007; Zhang et al., 2010) and the number of down-regulated genes belonging to this pathway, we tested the global impact of Bcl6 on the Wnt pathway *in vivo. Axin 2*, a classical Wnt/β-catenin-dependent target gene, was found to be up-regulated in *Bcl6^-/-^* mouse embryonic cortex using *in situ* hybridization. While *Axin2* expression is normally detected in the medial pallium of the frontal cortex in wild-type animals, a higher signal was found throughout dorsolateral levels in *Bcl6^-/-^* mice (Figure 2A-B), suggesting that β-catenin/Tcf activity is increased in the mutant cortex. Interestingly this difference was not detectable at more posterior levels (Figure 2A-B) in accordance with the frontal high occipital low graded *Bcl6* expression (Tiberi et al., 2012a). These data suggest that Bcl6 neurogenic function could depend on the down-regulation of the canonical Wnt pathway. We first tested this *in vitro* by examining potential genetic interactions between Bcl6 and β-catenin, the main signaling hub protein of the pathway. Neurogenic genes up-regulated *in vitro* by Bcl6 were prevented by CHIR99021, a GSK3 inhibitor over-activating the canonical Wnt pathway. CHIR99021, which increased the levels of the Wnt reporter gene *Lef1*, also prevented Bcl6-mediated down-regulation of Wnt target genes but did not prevent repression of Notch targets (Supplementary Figure S1). We then examined *in vivo* the impact of the gain of function of a stabilized β- catenin mutant on *Bcl6* overexpression using *in utero* electroporation (Figure 2). *Bcl6* gain of function led to an increased neurogenesis, as assessed by increased number of cells in the cortical plate (CP) and an increase in Tuj1+ neuronal cells at the expense of VZ-located-Pax6+ progenitors (Figure 2C-E). Importantly these effects were suppressed by overexpression of stabilized β-catenin.

**Figure 2.**
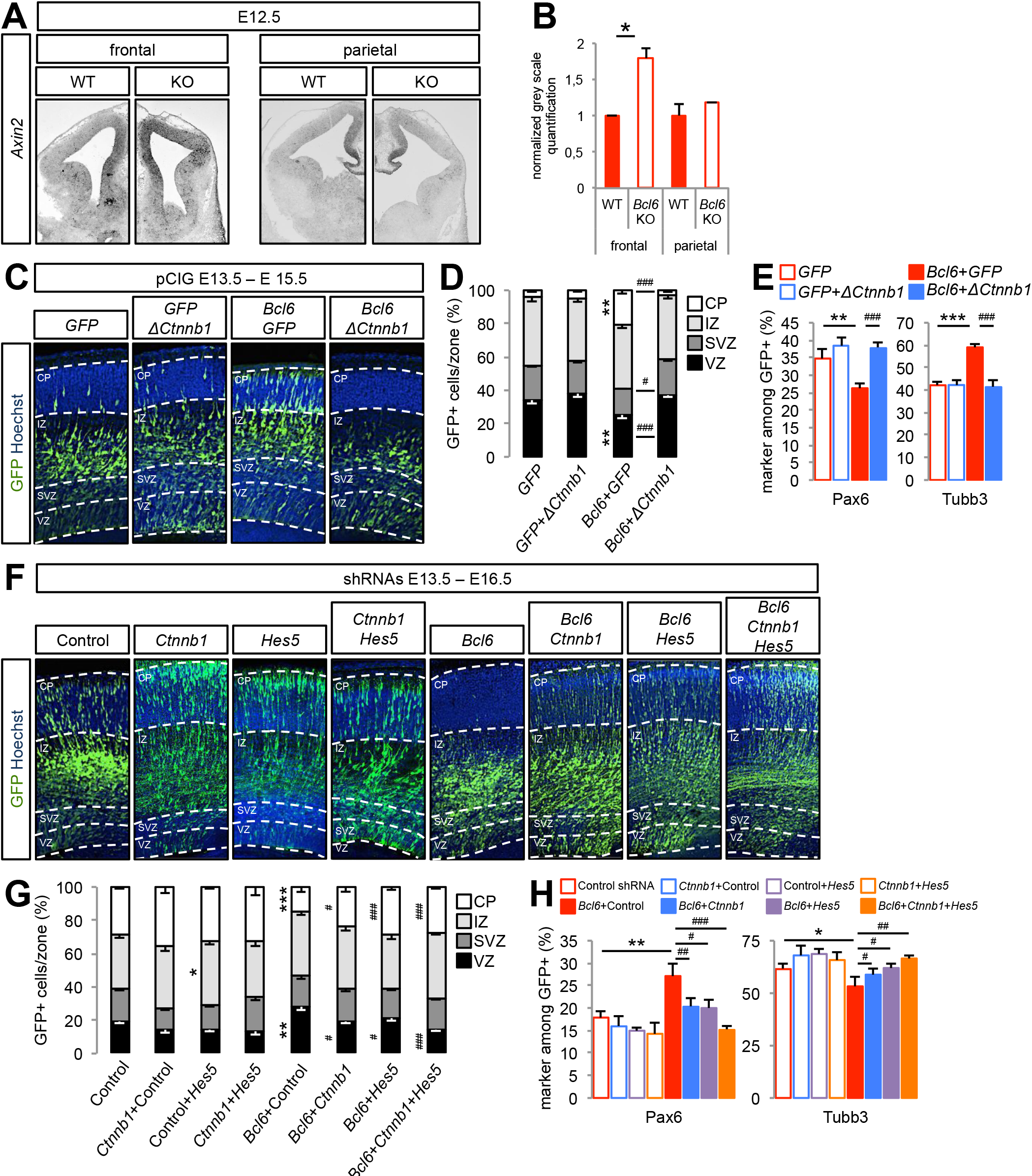
*Bcl6* alters β-catenin signaling *in vivo* to promote neurogenesis. (A) *In situ* hybridization of *Axin2* Wnt reporter gene on coronal sections of E12.5 wild-type and *Bcl6*-/- frontal and parietal telencephalon. (B) Normalized gray scale quantifications of *Axin2* levels were performed using Image J software. Data are presented as mean + s.e.m. ^*^ *P*<0.05 using Student’s *t-*test. (C-E) *In utero* electroporation of pCIG, pCIG+pCIG -Δ(1-90)*Ctnnb1*, pCIG-*Bcl6*+pCIG or pCIG-*Bcl6*+pCIG-Δ(1-90)*Ctnnb1* at E13.5. (C) Hoechst and GFP immunofluorescence was performed on coronal sections of E15.5 brains. Dashed lines mark the basal and apical margins of the ventricular zone (VZ), subventricular zone (SVZ), intermediate zone (IZ) and cortical plate (CP). (D) Histograms show the percentage of GFP+ cells in VZ, SVZ, IZ and CP. ^**^ *P*<0.01 pCIG-*Bcl6*+pCIG vs. pCIG and ^#^ *P*<0.05, ^###^ *P*<0.001 pCIG-*Bcl6*+ pCIG-Δ(1-90)*Ctnnb1* vs. pCIG-*Bcl6*+pCIG using two-way ANOVA followed by Tukey post-hoc test. (E) Histograms show the percentage of Pax6+ and Tuj1+ cells among the GFP+ cells. ^**^ *P*<0.01, ^***^ *P*<0.001 pCIG-*Bcl6*+pCIG vs. pCIG+pCIG and ^###^ *P*<0.001 pCIG-*Bcl6*+pCIG-Δ (1-90)*Ctnnb1* vs. pCIG-*Bcl6*+pCIG using one-way ANOVA followed by Tukey post-hoc test. Data are presented as mean + s.e.m. of n=6 control embryos (1856 cells), n=10 embryos for stabilized β-catenin gain of function (2727 cells), n=9 embryos for Bcl6 gain of function (1922 cells) and n=9 embryos for Bcl6 and stabilized β-catenin double gain of function (1588 cells). (F-H) *In utero* electroporation of scramble (control), scramble+*Ctnnb1*, scramble+*Hes5*, scramble+*Ctnnb1*+*Hes5*, scramble+*Bcl6*, scramble+*Ctnnb1*+*Bcl6*, scramble+*Hes5*+*Bcl6*, and *Ctnnb1*+*Hes5*+*Bcl6* shRNAs at E13.5. (F) Representative images of Hoechst and GFP immunofluorescence performed on coronal sections of E16.5 brains. Dashed lines mark the basal and apical margins of the ventricular zone (VZ), subventricular zone (SVZ), intermediate zone (IZ) and cortical plate (CP). (G) Histograms show the percentage of GFP+ cells in VZ, SVZ, IZ and CP. ^*^ *P*<0.05, ^**^ *P*<0.01, ^***^ *P*<0.001 vs. Control and ^#^ *P*<0.05, ^###^ *P*<0.001 vs. Bcl6+Control shRNAs using two-way ANOVA followed by Tukey post-hoc test. (H) Histograms show the percentage of Pax6+ and Tuj1+ cells among the GFP+ cells. ^*^ *P*<0.05, ^**^ *P*<0.01 vs. Control and ^#^ *P*<0.05, ^##^ *P*<0.01, ^###^ *P*<0.001 vs. Bcl6+Control shRNAs using one-way ANOVA followed by Tukey post-hoc test. Data are presented as mean + s.e.m. of n=15 control embryos (4476 cells), n=7 embryos for *Ctnnb1* shRNA (1851 cells), n=13 embryos for *Hes5* shRNA (3893 cells), n=7 embryos for *Ctnnb1*+*Hes5* shRNA (1932 cells), n=6 embryos for *Bcl6* shRNA (2093 cells), n=9 embryos for *Bcl6+Ctnnb1* shRNA (2642 cells), n=10 embryos for *Bcl6*+*Hes5* shRNA (2614 cells) and n=9 embryos for *Bcl6*+*Ctnnb1*+*Hes5* shRNA (2993 cells).

Conversely, we combined *Bcl6* and β-catenin (*Ctnnb1*) knockdown to assess potential epistasis. As expected (Tiberi et al., 2012a), *Bcl6* knockdown led to decreased neurogenesis, as assessed by increased cell number in the VZ at the expense of the CP (Figure 2F-G) associated with increased levels of Pax6+ progenitors and decreased Tuj1+ neurons (Figure 2H). This phenotype was significantly rescued by the *Ctnnb1* knock-down (Figure 2F-H). These data suggest that Bcl6 functionally acts on the Wnt pathway to promote neurogenesis, in addition to its effect on the Notch target *Hes5* (Tiberi et al., 2012a). We next directly compared the impact of Bcl6 on Wnt and Notch pathways *in vivo* by combining *Ctnnb1* and *Hes5* shRNAs to further assess whether Bcl6 represses them in parallel or sequentially. *Hes5* knockdown rescued some of the *Bcl6* loss of function-mediated phenotype to levels similar to the rescue obtained using the *Ctnnb1* knockdown. However, the association of both *Ctnnb1* and *Hes5* shRNA showed additive rescue of *Bcl6* knockdown reaching the corresponding control levels (*Ctnnb1*+*Hes5* shRNA combination; Figure 2F-H), suggesting that Bcl6 alters these two cascades at least in part in parallel.

Altogether these data indicate that Bcl6-mediated repression of the Wnt pathway, already at the level of β-catenin, is necessary to elicit neurogenic activity, in parallel to Notch signaling repression, and thus that Bcl6 acts by repression of multiple pathways promoting progenitor self-renewal and proliferation.

### Bcl6 promotes neurogenesis through Cyclin D inhibition

While Bcl6 appears to repress multiple serial components of individual pathways, it could also act through common effectors of parallel signals. In line with this hypothesis, *Cyclin d1/d2* genes were found to be down-regulated by Bcl6 overexpression *in vitro* (Figure 1D, Supplementary Table S1), and are known to be up-regulated by pathways driving progenitor self-renewal, including Wnt (Shtutman et al., 1999) but also SHH (Kasper et al., 2006; Katoh and Katoh, 2009), FGF/IGF (Kalita et al., 2013; Nilsson et al., 2012), and Notch (Cohen et al., 2010). Moreover, Cyclin d1/d2 are key promoters of cortical progenitor proliferation and consequently block neurogenesis (Lange et al., 2009; Pilaz et al., 2009; Tsunekawa et al., 2012).

We first examined their expression in the developing cortex in wild-type and Bcl6^-/-^ brains. *Ccnd1* was found in the VZ and SVZ in wild-type mice as previously reported (Glickstein et al., 2007), and its levels were significantly increased in *Bcl6^-/-^* animals (Figure 3A-B). *Ccnd2* mRNA was mostly detected in basal end-feet of radial glial cells and at lower intensity in the ventricular zone at E12.5, as previously described (Tsunekawa et al., 2012). While the high density of labeling and limited resolution of in situ hybridization precluded to detect upregulation in the basal end-feet, *Ccnd2* levels were significantly increased in the VZ in *Bcl6^-/-^* cortex (Figure 3C-D). Hence *Bcl6* negatively controls *Ccnd1/2* expression in cortical progenitors *in vivo*. We next examined the functional impact of *Bcl6* loss of function on *Ccnd1*/*Ccnd2* double knockdown as these two cyclins regulate cell-cycle progression in a redundant manner (Ciemerych et al., 2002; Glickstein et al., 2007; Tsunekawa et al., 2012). Decreased neurogenesis observed following *Bcl6* shRNA (Figure 3E-G) was completely rescued by the dual *Ccnd1*/*Ccnd2* knockdown (Figure 3E-G). This indicates that, in addition to repressing specific signaling pathway components, Bcl6 action also involves repression of common terminal effector targets driving progenitor proliferation and self-renewal.

**Figure 3.**
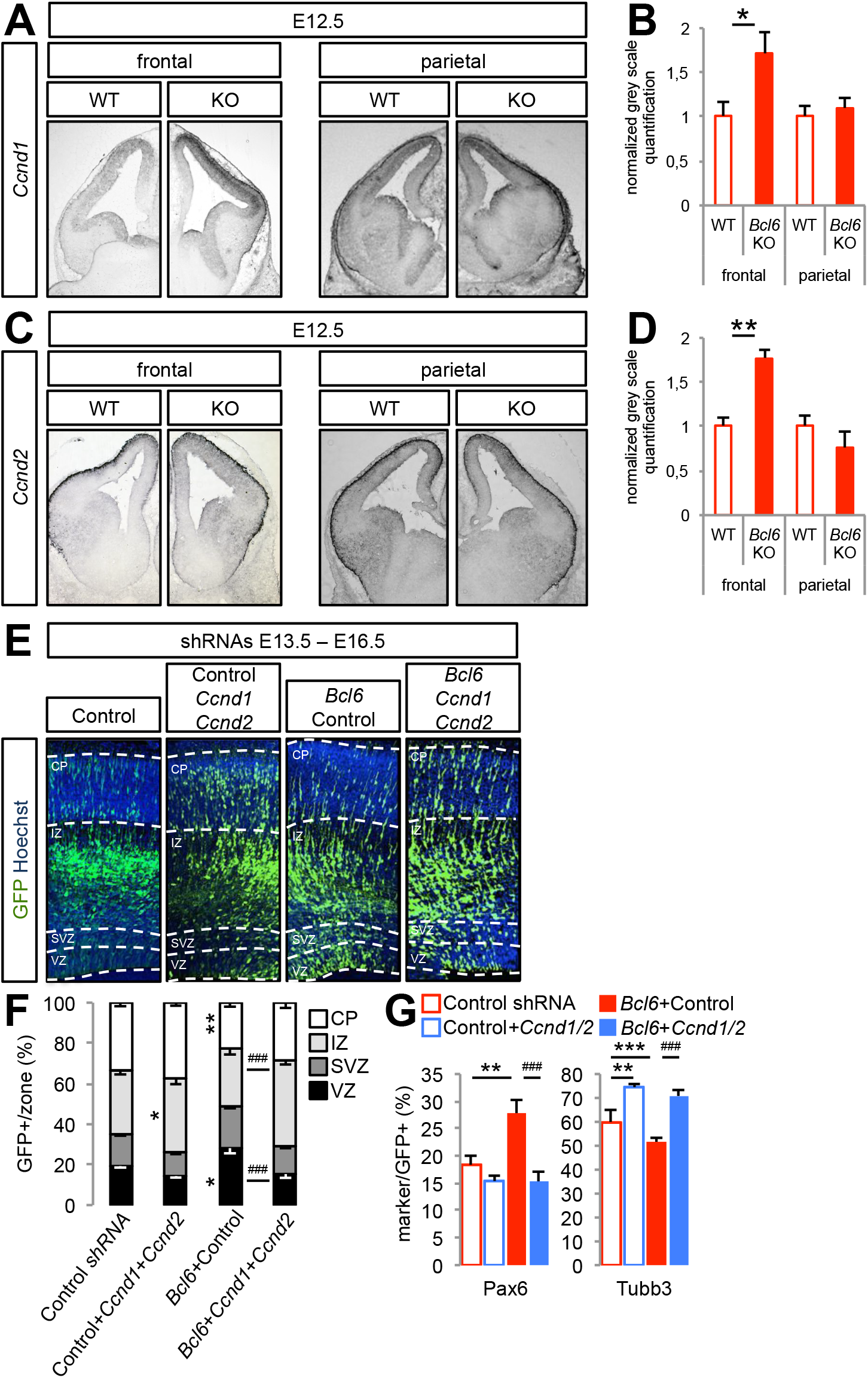
*Bcl6* directly affects *Ccnd1*/*Ccnd2* expression *in vivo* to promote neurogenesis. (A-D) *In situ* hybridization of (A) *Ccnd1* and (C) *Ccnd2* genes on coronal sections of E12.5 wild-type and *Bcl6*-/- frontal and parietal telencephalon. Normalized gray scale quantifications of (B) *Ccnd1* and (D) *Ccnd2* levels were performed using Image J software. Data are presented as mean + s.e.m. ^*^ *P*<0.05, ^**^ *P*<0.01 using Student’s *t-*test. (E-G) *In utero* electroporation of scramble (control), scramble+*Ccnd1*+*Ccnd2*, scramble+*Bcl6* and *Ccnd1*+*Ccnd2*+*Bcl6* shRNAs at E13.5. (E) Representative images of Hoechst and GFP immunofluorescence performed on coronal sections of E16.5 brains. Dashed lines mark the basal and apical margins of the ventricular zone (VZ), subventricular zone (SVZ), intermediate zone (IZ) and cortical plate (CP). (F) Histograms show the percentage of GFP+ cells in VZ, SVZ, IZ and CP. ^*^ *P*<0.01, ^**^ *P*<0.01 vs. Control and ^###^ *P*<0.001 vs. Bcl6+Control shRNAs using two-way ANOVA followed by Tukey post-hoc test. (G) Histograms show the percentage of Pax6+ and Tuj1+ cells among the GFP+ cells. ^**^ *P*<0.01, ^***^ *P*<0.001 vs. Control and ^###^ *P*<0.001 vs. Bcl6+Control shRNAs using one-way ANOVA followed by Tukey post-hoc test. Data are presented as mean + s.e.m. of n=12 control embryos (3627 cells), n=7 embryos for *Ccnd1*+*Ccnd2* shRNA (1519 cells), n=8 embryos for *Bcl6* shRNA (1998 cells) and n=12 embryos for *Bcl6*+*Ccnd1*+*Ccnd2* shRNA (3475 cells).

### Bcl6 down-regulates its target genes through generic as well as pathway-specific mechanisms

Overall our data indicate that Bcl6 acts through inhibition of multiple pathways to promote neurogenesis, raising the question of which effects are directly related to Bcl6 transcriptional repression, or reflect its indirect consequences. To address this point, we performed ChIP-seq to identify Bcl6 binding sites in *in vitro* cortical progenitors (Supplementary Figure S2). This revealed that Bcl6 binding was predominantly found on promoter regions, with a significant enrichment for Bcl6 matrix binding motif (Supplementary Figure S2A-D). Further, 39% of the Bcl6 predicted targets (1701/4366 peak-associated genes) were similar to those previously reported by ChIP-seq in human B cells (Basso et al., 2010). In line with our transcriptomics data, Bcl6 bound to numerous genes of the multiple cascades regulating the proliferation of cortical progenitors (Supplementary Figure S2E), although it should be noted that only 19% of the down-regulated genes from the RNA-seq dataset were found in the ChIP-seq screen (115/610 genes), indicating partial discrepancy between transcription factor binding and transcriptional effect, at least at the time points tested.

Then we validated *in vivo* some of these targets using ChIP-qPCR on the identified binding peaks in E12.5 wild-type and *Bcl6^-/-^* cortex, focusing on Wnt and Notch pathways. We first confirmed Bcl6 binding to the Notch target *Hes5*, but also identified additional Bcl6/Notch targets such as *Hes1* and *Nrarp* (Figure 4; Supplementary Figure S3A). Moreover, we found that Bcl6 bound to a repertoire of VZ-expressed genes of the Wnt/β-catenin cascade as well as potential Wnt/β-catenin targets, among which 10 showed significant down-regulation upon Bcl6 induction in *in vitro* ES cell-derived cortical progenitors, whether by RNA-seq or RT-qPCR, suggesting a direct functional impact following Bcl6 binding (Figure 4A, Supplementary Figure S4A). The genes bound and regulated by Bcl6 ranged from Wnt receptors/co-receptors to intracellular effectors to Wnt target genes (Figure 4B, Supplementary Figure S4B, Figure 1E).

**Figure 4.**
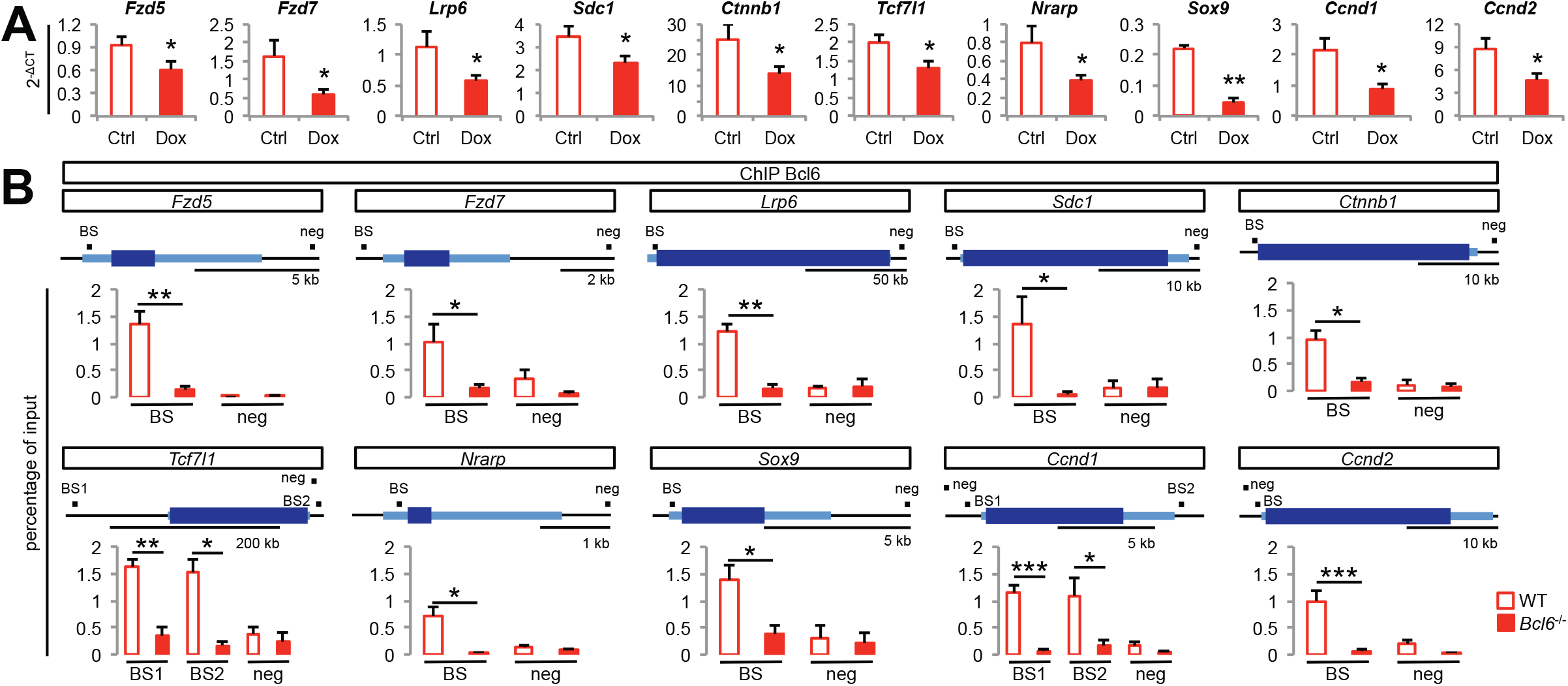
Bcl6 binds to core genes of the Wnt/β-catenin pathway to alter their expression. (A) RT-qPCR analysis of Wnt-related genes from DMSO- and doxycycline-treated cells at day 12. Data are presented as mean + s.e.m. of absolute levels (*n* = 21 from 8 differentiations). ^*^ *P*<0.05, ^**^ *P*<0.01 using Student’s *t-*test. (B) ChIP-qPCR validation of screened Bcl6-binding sites on regulatory regions and negative control sites of significantly down-regulated Wnt-related genes in E12.5 wild-type and *Bcl6*-/- telencephalon using a Bcl6 antibody. Data are presented as mean + s.e.m. of input enrichment (n = 4). ^*^ *P*<0.05, ^**^ *P*<0.01, and ^***^ *P*<0.001 using Student’s *t-*test. Genes are represented using 5’ to 3’ orientation with light blue showing 5’UTR and 3’UTR exons and thicker dark blue showing the exons of the coding sequence and the introns according to RefSeq gene sequences from mouse genome mm10 assembly (https://genome.ucsc.edu). Black boxes indicate amplified regions (BS: region comprising the Bcl6-predicted Binding Site inside the ChIP-seq significant peaks or NEG: region with no predicted Bcl6 matrix using the Jaspar software (http://jaspar.genereg.net)). See also Supplementary Figures S3 and S4.

We next examined how Bcl6 binding to its target sequences might lead to transcriptional repression. We previously found that Bcl6-mediated *Hes5* repression occurs through modifications of histone acetylation mediated by Sirt1 deacetylase recruitment (Tiberi et al., 2012a). Remarkably we found a similar mechanism for all tested target genes identified above (Figure 5). Indeed, *in vivo* ChIP-qPCR experiments performed on mouse E12.5 embryonic cortex showed enrichment for acetylated lysines H4K16 and H1.4K26 on all tested genes in *Bcl6^-/-^* cortex compared with wild-type, indicating that repressive chromatin remodeling required the presence of Bcl6 binding (Figure 5A, Supplementary Figure S3B). Moreover, we detected binding of the histone deacetylase Sirt1 to all Bcl6-bound target genes, which was abolished in *Bcl6^-/-^* cortex (Figure 5B, Supplementary Figure S3C).

**Figure 5.**
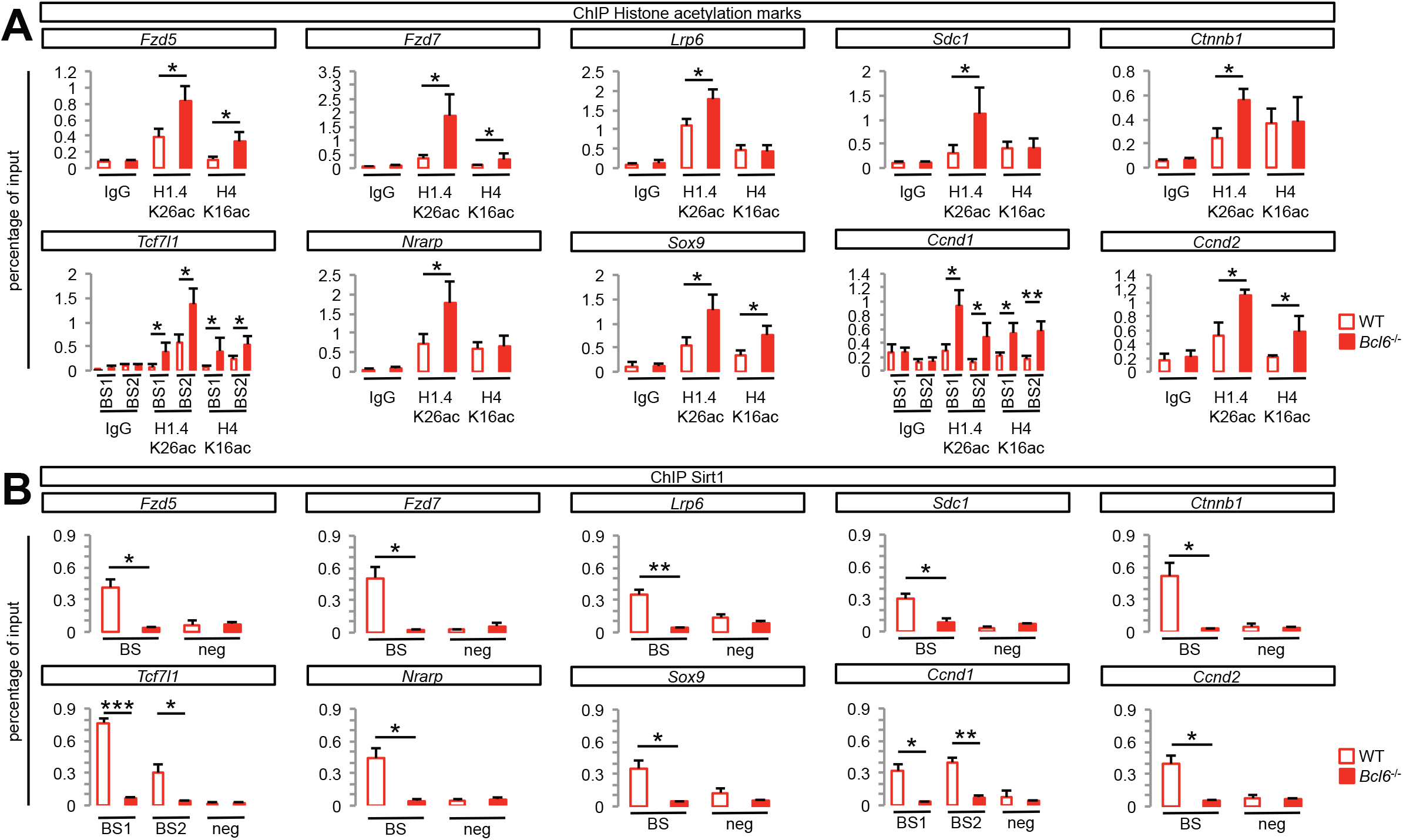
Bcl6 binding to significantly down-regulated Wnt-related genes induces chromatin remodeling and Sirt1 recruitment. (A) ChIP-qPCR of histone acetylation marks on Bcl6 binding sites of significantly down-regulated Wnt-related genes in E12.5 wild-type and *Bcl6*-/- telencephalon using control (rabbit IgG), H1.4K26ac and H4K16ac antibodies. Data are presented as mean + s.e.m. of input enrichment (n = 6). ^*^ *P*<0.05 and ^**^ *P*<0.01 using Student’s *t-*test. (B) ChIP-qPCR on validated Bcl6 binding sites of significantly down-regulated Wnt-related genes in E12.5 wild-type and *Bcl6*-/- telencephalon using a Sirt1 antibody. Data are presented as mean + s.e.m. of input enrichment (n = 6). ^*^ *P*<0.05, ^**^ *P*<0.01, and ^***^ *P*<0.001 using Student’s *t-*test. See also Supplementary Figure S3.

These data point to a generic mechanism by which a wide repertoire of the genes down-regulated by Bcl6 and belonging to stem cell renewal signaling pathways are directly bound and repressed by Bcl6, which then recruits Sirt1, leading to their transcriptional silencing.

To test whether additional mechanisms that are signaling pathway-specific may be involved, we examined in more depth the mechanism of action of Bcl6 on *Ccnd1/2.* Bcl6 was found to bind to their regulatory regions *in vitro*, together with Sirt1, in association with histone H4K16 deacetylation, as observed *in vivo* (Figure 6). We then examined the binding profile of Tcf7l1 (Figure 6A-B), the only Wnt/β-catenin-associated Tcf/Lef transcription factor known to be expressed in cortical progenitors (Galceran et al., 2000). Tcf7l1 was found to bind to both *Ccnd1/2* promoters on predicted Tcf/Lef binding sites, confirming their direct link with the Wnt pathway (Figure 6B). Most strikingly, this binding was decreased upon Bcl6 overexpression (Figure 6B). In contrast, Tcf7l1 binding at the level of the *Lef1* promoter, to which Bcl6 does not bind to, was unaffected (Supplementary Figure S5), suggesting that Bcl6 directly affects Tcf7l1 binding to *Ccnd1/2* promoters.

**Figure 6.**
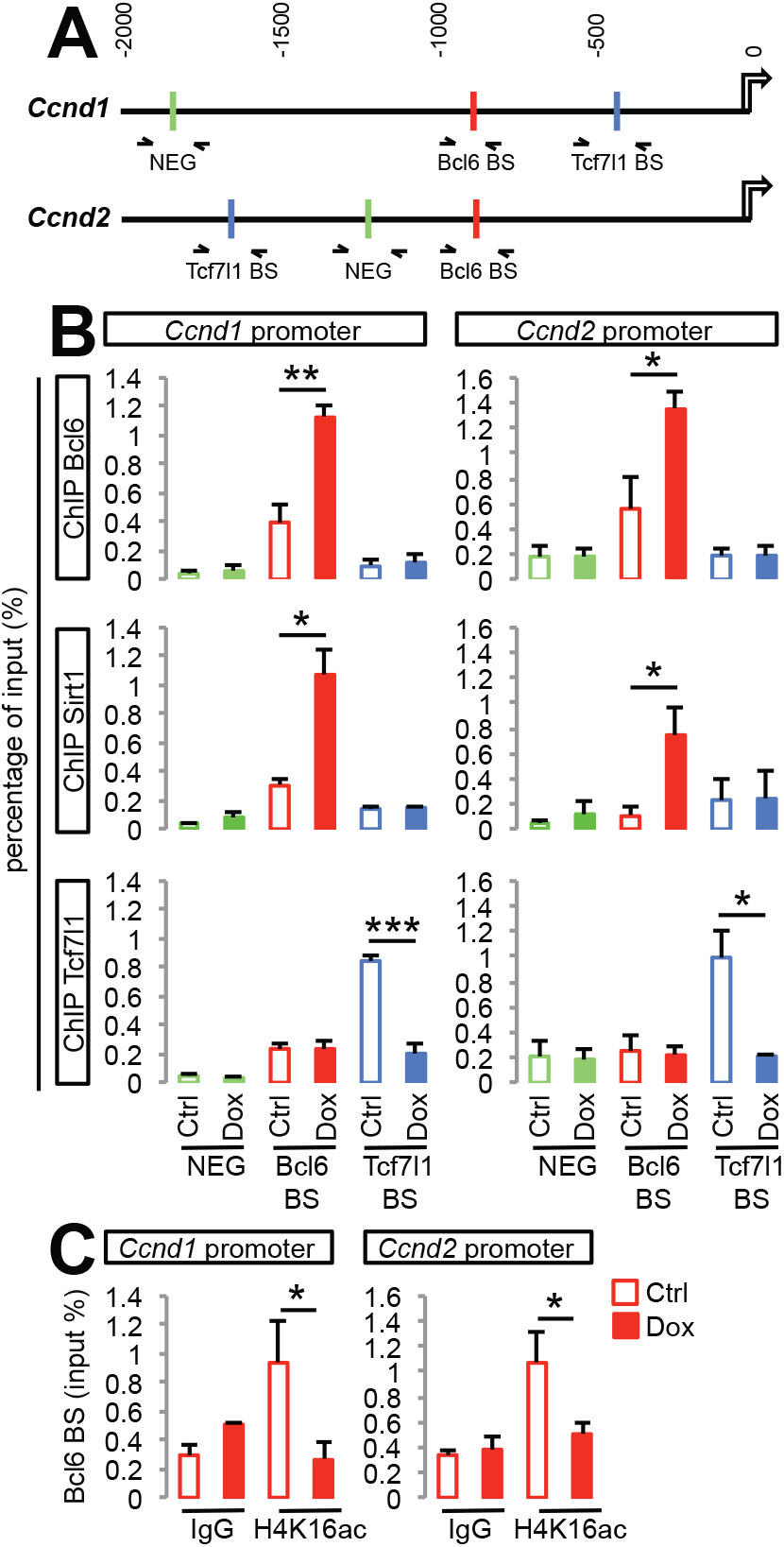
Bcl6 and Sirt1 bind to *Ccnd1* and *Ccnd2* regulatory regions leading to the removal of the β-catenin effector Tcf7l1. (A) Schematic representation of the genomic region 2 kb upstream from *Ccnd1* and *Ccnd2* transcription starting sites showing validated Bcl6 (red) as well as putative Tcf7l1 (blue) binding sites and negative regions for either transcription factor (green) as predicted by the Jaspar software (http://jaspar.genereg.net). The arrows represent the amplified regions by qPCR used to measure the enrichment following ChIP. (B) ChIP-qPCR analysis of the Bcl6, Sirt1 and Tcf7l1 binding sites on the *Ccnd1* and *Ccnd2* regulatory regions in cortical progenitors derived from *Bcl6* A2 lox.Cre mouse ES cells (differentiation day 12, 24h DMSO (Ctrl) or Dox treatment). Data are presented as mean + s.e.m. of input enrichment (n = 3 differentiations). ^*^ *P*<0.05, ^**^ *P*<0.01, ^***^ *P*<0.001 using Student’s *t-*test. (C) ChIP-qPCR analysis of the H4K16ac histone acetylation mark on Bcl6 binding sites of the *Ccnd1* and *Ccnd2* regulatory regions in cortical progenitors derived from *Bcl6* A2 lox.Cre mouse ES cells (differentiation day 12, 24h DMSO (Ctrl) or Dox treatment). Data are presented as mean + s.e.m. of input enrichment (n = 3 differentiations). ^*^ *P*<0.05 using Student’s *t-*test. See also Supplementary Figure 5.

These data suggest that Bcl6 has specific effects on the binding of other transcription factors to promoters, as an additional mechanism to modulate responsiveness to extrinsic pathways.

## Discussion

Neurogenesis is a key fate transition controlled by the interplay of intrinsic and extrinsic cues, but how these are integrated at the molecular level remains poorly known. Here we show how a single transcriptional repressor, Bcl6, can promote fate conversion through the direct repression of a repertoire of genes encoding key components of numerous extrinsic morphogen pathways that promote self-renewal and proliferation. This provides important insight on the molecular logic by which neurogenic conversion can occur in a robust fashion in the presence of many, and sometimes contradictory, extrinsic cues. Our transcriptome analysis shows that Bcl6 gain of function elicits a broad neurogenic program, from genes involved in the generation and expansion of intermediate progenitors, their differentiation into neurons, as well as neuronal maturation and specification, similarly to the response to the proneural *Neurog2* transcription factor in the developing cortex (Gohlke et al., 2008; Johnson et al., 2015). However, in the case of Bcl6, these effects are indirectly linked to a direct negative impact of Bcl6 on pathways promoting self-renewal and proliferation. Indeed, Bcl6 represses the transcription of genes related to most signaling cascades that favor the proliferation of cortical progenitors at the expense of their differentiation, most strikingly the Notch and Wnt pathways, as well as SHH and FGF. Combining our RNA-seq results with ChIP experiments, we found that Bcl6 directly binds to many of these genes leading to their transcriptional repression, even if future work will be needed to describe in depth Bcl6 direct repression on pathways other than Notch and Wnt.

This global repressive effect is reminiscent of the one previously proposed for Myt1l in the frame of neuronal reprogramming, where a single transcription factor appears to repress globally multiple non-neuronal fates to promote *in vitro* conversion into neurons (Mall et al., 2017) or of Lola and Prospero that repress neural stem cell fate in fly neurons (Southall et al., 2014). It is also strikingly complementary to the Rest complex, which represses neuronal fate genes in non-neuronal cells (Ballas et al., 2005). In the case of Bcl6 however, our data suggest that its repression does not appear to act so much on fate-specific genes *per se*, but rather on genes selectively involved in responsiveness to extrinsic signaling pathways promoting progenitor maintenance. In this sense Bcl6 acts probably mostly during the fate transition itself, and could provide a mechanism for the progressive restriction from extrinsic to intrinsic cues that has been long described following neuronal differentiation (Edlund and Jessell, 1999), in line with the described expression pattern of Bcl6 that gets gradually upregulated as neural progenitors undergo neuronal differentiation (Tiberi et al., 2014; Tiberi et al., 2012a). This molecular logic could also partially ensure that neuronal differentiation takes place irreversibly, even in the presence of proliferative extrinsic cues, thus providing robustness to the neurogenesis process. In the same frame, as Bcl6 was previously found to act as an oncosuppressor in the cerebellum (Tiberi et al., 2014), it will be interesting to test whether this effect may be linked to a block of dedifferentiation in this or other neural cell types, as shown for the transcriptional repressor Lola in the fly nervous system (Southall et al., 2014).

Interestingly in the case of Wnt signaling, Bcl6 appears to act upon multiple components, from receptors to transducers and transcriptional effectors, as attested by the functional interactions between Bcl6 and β-catenin, as well as with Ccdn1/2, revealed in this study. This implies that while Bcl6 can alter the intracellular response of Notch signaling in a target-specific way (Sakano et al., 2010; Tiberi et al., 2012a), its effect on Wnt signaling appears to be much more global, from responsiveness to output. This may be related to the high complexity of the Wnt pathway, such as multiplicity of ligands and receptors and cell-dependent positive and negative feedback loops (Clevers and Nusse, 2012; MacDonald et al., 2009), on which multiple levels of repression may be required to achieve robust silencing. Indeed it should be noted that although many of the Wnt components that are repressed are positive regulators of the pathway, some inhibitors of the pathway, such as *Sfrp1/2*, were also downregulated, while some activators such as *Fzd8* or inhibitors such as *Dkk1* were, most likely indirectly, upregulated. The same is true for the other examined morphogen pathways, thus suggesting that, although the overall effect of Bcl6 on these pathways is inhibitory, the effect on individual genes could work in both directions.

Moreover, the comparison of combined epistasis of *Bcl6* with *Hes5* and *Ctnnb1* revealed that Bcl6 appears to affect Notch and Wnt signaling in a parallel fashion rather than through a cascade of events following the alteration of a single upstream pathway. Furthermore, some Bcl6-elicited gene expression changes impact several pathways at the same time. For instance, we observed that Bcl6 gain of function decreases the expression of *Jag1* and *Nrarp*, which are both linked to Notch but also Wnt pathways (Ishitani et al., 2005; Phng et al., 2009).

We identify *Ccnd1* and *Ccnd2* as major key targets of Bcl6, and their down-regulation rescues to a large extent the loss of *Bcl6*. *Ccnd1* or *Ccnd2* were previously shown to act as key promoters of cortical progenitor proliferation and blockers of differentiation (Lange et al., 2009; Pilaz et al., 2009; Tsunekawa et al., 2012) and are found downstream of most signaling cues promoting progenitor self-renewal. This suggests a model whereby Bcl6 contributes to robustness of fate transition by also acting on key effector targets lying downstream to several pathways. In addition, the regulation of *Ccnd1* and *Ccnd2* by Bcl6 could contribute to other aspects of cortical development such as area patterning, given the enriched expression of Bcl6 in the frontal cortex (Tiberi et al., 2012a). Indeed, it has been suggested that the specification of the cortex topology could rely in part on rostro-caudal and latero-medial gradients of the cell-cycle G1 phase duration and neurogenesis (Miyama et al., 1997), to which Bcl6 could contribute to in a major way, given its preferential expression in the frontal cortex, and its effects on signaling cues, such as Wnts and FGFs that can also impact on cortical areal patterning (Sur and Rubenstein, 2005; Tiberi et al., 2012a).

From a molecular viewpoint, Bcl6 appears to act through the same generic mechanism on most if not all of its targets, i.e. direct repression and Sirt1 recruitment, together with histone deacetylation that is correlated with decreased transcription. However some aspects of Bcl6 function are also pathway-specific, such as the regulation of Tcf7I1 on *Ccnd1/2* genes. It will be interesting to determine the underlying precise mechanism, and whether Bcl6 regulates other pathway-specific transcription factors in a similar way.

To sum up, our data support a model whereby a single intrinsic factor downregulates the responsiveness to extrinsic cues, through transcriptional repression at multiple parallel and serial levels along the downstream pathways, to ensure irreversible neurogenic fate transition. Our data thus uncover a novel and elegant mechanism that controls the neurogenic transition, and establish an important molecular link between transcriptional repression and robustness of neural fate conversion. As Bcl6 is expressed in only specific subsets of progenitors and neurons during brain development, future work should determine whether other transcriptional repressors in distinct parts of the nervous system can similarly contribute to neurogenic fate transition.

## Acknowledgments

The authors thank G. Vassart for continuous support and interest, members of the Vanderhaeghen lab and Institute for Interdisciplinary Research for helpful discussions and advice, J.-M. Vanderwinden of the Light Microscopy Facility for his support with imaging, F. Libert and A. Lefort (Brussels Interuniversity Genomics High Throughput core) and D. Gacquer for RNA-seq, The Advanced Sequencing Team at the Francis Crick Institute for ChIP-seq, B. Merrill (University of Illinois) for kindly sharing Tcf7l1 antibody, and R. Dalla-Favera (Columbia University) for generously sharing the *Bcl6-/-* mice. This work was funded by the Belgian FRS/FNRS, the European Research Council (ERC Adv Grant GENDEVOCORTEX), the FMRE, the Interuniversity Attraction Poles Program (IUAP), the WELBIO Program of the Walloon Region, the AXA Research Fund, the Fondation ULB, the ERA-net ‘Microkin’ (to PV), and the Vlaams Instituut voor Biotechnologie (VIB) (to PV and SA). The work done in FG’s lab was funded by the Francis Crick Institute, which receives its core funding from Cancer Research UK (FC0010089), the UK Medical Research Council (FC0010089), and the Wellcome Trust (FC0010089).

## Author Contributions

Conceptualization and Methodology, J.B., L.T., J.v.d.A., and P.V.; Investigation, J.B. and L.T. with the help of J.v.d.A., Z.B.G., A.B., A.H. and F.D.V.B.; Formal Analysis, J.B., D.P. and X.L.; Writing - Original Draft, J.B. and P.V.; Writing - Review & Editing, J.B., L.T., J.v.d.A., D.P., F.G., S.A. and P.V.; Funding acquisition, P.V.; Resources, F.G., S.A. and P.V.; Supervision, F.G., S.A. and P.V.

## STAR Methods

### Animals

All mouse experiments were performed with the approval of the Université Libre de Bruxelles Committee for animal welfare. Animals were housed under standard conditions (12 h light:12 h dark cycles) with food and water *ad libitum*. For *in utero* electroporation experiments, timed-pregnant mice were obtained by mating adult RjOrl:SWISS CD1 mice (Janvier, France). The plug date was defined as embryonic day E0.5. For experiments using *Bcl6^-/-^* mice, we mated *Bcl6*+/− mice (mixed C57BL/6 and CD1 background), in which the null allele lacked exons 4–10 (Ye et al., 1997).

### Plasmids

The *Ccnd1* coding sequences (888 bp) and the *Ccnd2* 5’UTR+coding sequence (1145 bp) were amplified by PCR from ESC-derived cortical progenitor cDNA, and the sequences verified and cloned into pGEMT plasmid (Promega). The *Axin2* plasmid was a kind gift from Dr Isabelle Garcia (ULB).

The *Bcl6* coding sequence was amplified by PCR from cDNA and cloned into p*CAG*-IRES-*GFP* (pCIG). The p*CAG*- Δ 90*Ctnnb1* plasmid (N-terminal deletion lacking GSK3-dependent S^33^, S^37^ and T^41^ phosphorylation sites) was obtained from Addgene (plasmid #26645). shRNA plasmids were cloned downstream of the *U6* promoter into the pSilencer2.1-CAG-Venus (pSCV2)- plasmid as previously described (Tiberi et al., 2012a) with the exception of the *Ctnnb1* shRNA plasmid (pLKO.1-puro vector, Sigma). Target sequences were 5’-ACTACCGTTGTTATAGGTG-3’ (Control; (Tiberi et al., 2012a), 5’- TGATGTTCTTCTCAACCTTAA-3’ (*Bcl6*; (Tiberi et al., 2012a; Zhang et al., 2007), 5’-CCCAAGCCTTAGTAAACATAA-3’ (*Ctnnb1*; (Mao et al., 2009), 5’- AGCCTGCACCAGGACTAC-3’ (*Hes5*; (Lee et al., 2007), 5’- CCACAGATGTGAAGTTCATTT-3’ (*Ccnd1*; (Jirawatnotai et al., 2011), 5’- CGACTTCAAGTTTGCCATGTA-3’ (*Ccnd2*; (Gargiulo et al., 2013) and were previously validated.

### In Utero Electroporation

Timed-pregnant mice were anesthetized with a ketamine/xylazine mixture at E13.5. Each uterus was cautiously exposed under sterile conditions. Fast-green (Sigma)-labeled plasmid solutions were prepared using either 1 μg/μL or 0.5 μg/μL of appropriate DNA combinations for gain of function experiments or knockdown experiments, respectively. DNA solutions were injected into the lateral ventricles of the embryos using a heat-pulled glass capillary (1.0 OD × 0.78 × 100 L mm; Harvard Apparatus) prepared with a heat micropipette puller (heat: 580, pull: 100, velocity: 170, time: 120, pressure: 500; Sutter Instrument P-87). Electroporation was performed using tweezers electrodes (Nepa Gene CUY650P5) connected to a BTX830 electroporator (5 pulses of 30 V for 100 ms with an interval of 1 s). Embryos were placed back into the abdominal cavity, and mice were sutured and placed on a heating plate until recovery.

### Immunofluorescence

Embryos were fixed by transcardiac perfusion with freshly-prepared 4% paraformaldehyde (Invitrogen). Brains were dissected, and 100 mm sections were prepared using a Leica VT1000S vibrosector. Slices were transferred into PBS with 0.5 μg/mL sodium azide (Sigma), then blocked with PBS supplemented with 3% horse serum (Invitrogen) and 0.3% Triton X-100 (Sigma) during 1 h, and incubated overnight at 4°C with the following primary antibodies: chicken GFP (Abcam, 1:2,000), rabbit Pax6 (Covance, 1:1000), mouse β3-tubulin (Tuj1 epitope, Covance, 1:1000), or rabbit Ki67 (Abcam, 1:500). After three washes with PBS/0.1% Triton X-100, slices were incubated in PBS for 1 h at room temperature and incubated 2 h at room temperature with the appropriated Alexa-488 (1:1,000, Molecular Probes) and Cyanine 3 (1:400, Jackson ImmunoResearch) secondary antibodies. Sections were again washed three times with PBS/0.1% Triton X-100, stained with Hoechst (bisBenzimide H 33258, Sigma) for 5 min and washed twice in PBS. The sections were next mounted on a Superfrost slide (Thermo Scientific) and dried using a brush before adding Glycergel mounting medium (Dako). Imaging was performed using a Zeiss LSM780 confocal microscope controlled by the Zen Black software (Zeiss).

### Mouse ES cells and cortical differentiation

The A2 lox.Cre Bcl6 cell lines, tetracyclin-inducible Bcl6 ICE (A2lox.Cre) mouse Embryonic Stem Cells (Tiberi et al., 2012a), were routinely propagated on irradiated mouse embryonic fibroblasts in DMEM (Invitrogen) supplemented with 15% ESC-certified fetal bovine serum (vol/vol, Invitrogen), 0.1 mM non-essential amino acids (Invitrogen), 1 mM sodium pyruvate (Invitrogen), 0.1 mM β- mercaptoethanol (Sigma), 50 U.ml^−1^ penicillin/ streptomycin and 103 U.ml^−1^ mouse Leukemia Inhibitor Factor (ESGRO). Results were obtained using three independent clones.

For differentiation, A2 lox.Cre Bcl6 mouse ESCs were plated at low density (20 × 103 ml^−^1) on gelatin-coated coverslips and cultured as previously described (Gaspard et al., 2009). Briefly, on day 0 of the differentiation, the ES medium was changed to DDM medium. At day 2, DDM was supplemented with cyclopamine (400 ng/mL, Calbiochem) and the medium was replenished every 2 days. After 10 days, medium was switched back to DDM. Doxycline treatment (1 μg.ml^−1^) for Bcl6 induction or DMSO (1:1000, control) was applied for 12-24 h at day 12. CHIR99021 (4 μM, Tocris) needed to be applied for 48h, i.e. from day 10, to effectively impact Wnt target genes.

### Transcriptome analyses

Total mRNA from four independent samples from ESc-derived cortical differentiations (day 12 ± doxycycline) was extracted using the QIAGEN RNeasy mini kit according to the manufacturer’s recommendations. Following RNA quality control assessed on a Bioanalyzer 2100 (Agilent technologies), the indexed cDNA libraries were prepared using the TruSeq stranded mRNA Library Preparation kit (Illumina) following manufacturer’s recommendations. The multiplexed libraries (18 pM) were loaded on flow cells and sequences were produced using a HiSeq PE Cluster Kit v4 and TruSeq SBS Kit v3-HS from a Hiseq 1500 (Illumina). Approximately 20 million of paired-end reads per sample were mapped against the mouse reference genome (GRCm38.p4/mm10) using STAR 2.5.3a software (Dobin et al., 2013) to generate read alignments for each sample. Annotations Mus_musculus.GRCm38.87.gtf were obtained from ftp.Ensembl.org. After transcript assembling, gene level counts were obtained using HTSeq 0.9.1 (Anders et al., 2015). EdgeR 3.20.1 (Robinson et al., 2010) was then used to calculate the level of differential gene expression. Gene Ontology analyses of biological processes were performed using the GOrilla application (http://cbl-gorilla.cs.technion.ac.il).

### RT-qPCR

Reverse transcription of mature mRNAs was done with the Superscript II kit (Invitrogen) using the manufacturer’s protocol for oligo dTs. Quantitative PCR (qPCR) was performed in duplicate using Power Sybr Green Mix (Applied Biosystems) and a 7500 Real-Time PCR System (Applied Biosystems). Results are presented as linearized Ct values normalized to the housekeeping gene *Tbp* (2-ΔCt) and the primers are listed in the Supplementary Table S4.

### *In situ* RNA hybridization

In situ hybridization was performed using digoxigenin-labeled RNA probes (DIG RNA labeling kit, Roche) as previously described (Lambot et al., 2005). The above-mentioned plasmids were linearized and reverse-transcribed using NotI and T3 (*Axin2*), SpeI and T7 (*Ccnd1*), and SacII and Sp6 (*Ccnd2*) to generate the anti-sense probes. Sense probes were used as a negative control for each gene tested and revealed no specific staining (data not shown). Littermate embryos were compared using 3 animals/genotype from 3 different litters for *Axin2* and *Ccnd2* or 6 embryos/genotype from 4 different litters (*Ccnd1*). Images were acquired with a Zeiss Axioplan 2 microscope and a Spot RT3 camera using the Spot 5.2 software. Quantifications of gray densities were done on comparable levels of the dorsal primordium of the frontal cortex using ImageJ software (Schindelin et al., 2012).

### Chromatin Immunoprecipitation

A2 lox.Cre Bcl6 cells and E12.5 *Bcl6* WT and KO telencephalons were fixed with 1% formaldehyde in phosphate-buffered saline and then lysed, sonicated, and immunoprecipitated as described previously (Castro et al., 2011). Immunoprecipitations were performed using IgG isotype control (ab171870, Abcam), Bcl6 (sc-858, Santa Cruz Biotechnology), H1.4K26ac (H7789, Sigma-Aldrich), H4K16ac (07-329, Millipore), Sirt1 (07-131, Millipore), and Tcf7l1 (kind gift from Prof. Brad Merrill) rabbit antibodies and 50/50 mixed protein A and protein G magnetic Dynabeads (ThermoFisher Scientific). Primers used for ChIP-qPCR were designed to surround the predicted Bcl6 matrices present in the significant peaks of the ChIP-seq screen and are listed in the Supplementary Table S4.

### ChIP-seq

Following chromatin immunoprecipitation using A2 lox.Cre Bcl6 cells (differentiation day 12, 24h Bcl6 induction using 1 μg.ml^−1^ Doxycline treatment) as described above, ChIP libraries were prepared according to the standard Illumina ChIP-seq protocol and were sequenced with the Genome Analyzer IIx (Illumina) as previously described (Mateo et al., 2015). Reads from three independent ChIP-seq experiments were pooled and aligned to mm10 mouse genome assembly using Bowtie (Langmead et al., 2009). Peaks were then called using MACS (Zhang et al., 2008) with standard parameters with default cutoff *P* value <10e-5. Target genes associated with peaks were identified by GREAT version 3.0 (McLean et al., 2010). ChIP-seq dataset for the Bcl6-predicted target genes were visualized using the UCSC Genome Browser (Kent et al., 2002). Search for an enrichment of Bcl6 matrices in the MACS-called ChIP peaks was performed using i-cisTarget (Imrichova et al., 2015) using a promoter-only database.

### Statistical analysis

Results are shown as mean^±^standard error (S.E.M.) of at least three biologically independent experiments. Student’s unpaired *t*-test was used for two group comparisons. Analyses of multiple groups were performed by a one-way or two-way analysis of variance (ANOVA) - as indicated in figure legends - followed by post hoc multiple comparisons using Tukey’s test. Hypothesis testing for overlap analysis was performed using the Genomic HyperBrowser (https://hyperbrowser.uio.no/(Sandve et al., 2010). For all tests, a *P*-value inferior to 0.05 was taken as statistically significant.

### Data and Software Availability

RNA-seq raw data are shown on Supplementary Table S5. ChIP-seq data can be found on the UCSC Genome Browser using the following Hyperlink : <u>UCSC display for ChIP-seq dataset</u>.

## Supplemental Information

**Supplementary Figures S1-S5.**

**Supplementary Table S1. Related to Figure 1. RNA-seq analysis upon Bcl6 induction in *in vitro* cortical progenitors.** RNA-seq dataset of the significantly up- and down-regulated genes following Bcl6 expression in cortical progenitors derived from A2 lox.Cre cells (differentiation day 12, 24 h DMSO (control) or doxycycline treatment).

**Supplementary Table S2. Related to Figure 1B. Gene Ontology analyses of the ranked up- and down-regulated genes from the RNA-seq dataset**. GO analysis was performed using the GOrilla software (http://cbl-gorilla.cs.technion.ac.il).

**Supplementary Table S3. Related to Figure 1D. Complete gene set used for the analysis of transcriptional changes elicited by Bcl6 overexpression on Wnt, Notch, SHH, and FGF/insulin/IGF proliferative pathways in *in vitro* cortical progenitors.** Significantly up- and down-regulated genes are shown in red and blue, respectively. Genes marked with an asterisk are target genes of the pathway itself. Potential target genes highlighted in green show expression in cortical progenitors according to literature data when available or online data sets (http://developingmouse.brain-map.org, http://www.genepaint.org, http://rakiclab.med.yale.edu/transcriptome/).

**Supplementary Table S4. Oligonucleotide sequence for RT-qPCR and ChIP-qPCR.**

**Supplementary Table S5. RNA-seq raw data counts.**

